# Global network structure and local transcriptomic vulnerability shape atrophy in sporadic and genetic behavioral variant frontotemporal dementia

**DOI:** 10.1101/2021.08.24.457538

**Authors:** Golia Shafiei, Vincent Bazinet, Mahsa Dadar, Ana L. Manera, D. Louis Collins, Alain Dagher, Barbara Borroni, Raquel Sanchez-Valle, Fermin Moreno, Robert Laforce, Caroline Graff, Matthis Synofzik, Daniela Galimberti, James B. Rowe, Mario Masellis, Maria Carmela Tartaglia, Elizabeth Finger, Rik Vandenberghe, Alexandre de Mendonça, Fabrizio Tagliavini, Isabel Santana, Chris Butler, Alex Gerhard, Adrian Danek, Johannes Levin, Markus Otto, Sandro Sorbi, Lize C. Jiskoot, Harro Seelaar, John C. van Swieten, Jonathan D. Rohrer, Bratislav Mišić, Simon Ducharme, Frontotemporal Lobar Degeneration Neuroimaging Initiative (FTLDNI), GENetic Frontotemporal dementia Initiative (GENFI)

## Abstract

Connections among brain regions allow pathological perturbations to spread from a single source region to multiple regions. Patterns of neurodegeneration in multiple diseases, including behavioral variant of frontotemporal dementia (bvFTD), resemble the large-scale functional systems, but how bvFTD-related atrophy patterns relate to structural network organization remains unknown. Here we investigate whether neurodegeneration patterns in sporadic and genetic bvFTD are conditioned by connectome architecture. Regional atrophy patterns were estimated in both genetic bvFTD (75 patients, 247 controls) and sporadic bvFTD (70 patients, 123 controls). We first identify distributed atrophy patterns in bvFTD, mainly targeting areas associated with the limbic intrinsic network and insular cytoarchitectonic class. Regional atrophy was significantly correlated with atrophy of structurally- and functionally-connected neighbors, demonstrating that network structure shapes atrophy patterns. The anterior insula was identified as the predominant group epicenter of brain atrophy using data-driven and simulation-based methods, with some secondary regions in frontal ventromedial and anteromedial temporal areas. Finally, we find that FTD-related genes, namely C9orf72 and TARDBP, confer local transcriptomic vulnerability to the disease, effectively modulating the propagation of pathology through the connectome. Collectively, our results demonstrate that atrophy patterns in sporadic and genetic bvFTD are jointly shaped by global connectome architecture and local transcriptomic vulnerability.

## Introduction

Frontotemporal dementia (FTD) is one of the most common forms of early-onset dementia [39, 47]. The behavioral variant of FTD (bvFTD), which presents with various combinations of behavioral (apathy, disinhibition, compulsions and stereotypies), personality (decreased empathy and sympathy, altered personal preferences) and cognitive (executive dysfunction and social cognitive deficits) changes, is the most common clinical variant of FTD [39, 46]. Despite its distinctive clinical presentation, bvFTD is pathologically heterogenous, with the most common subtypes being related to the accumulation of hyperphosphorylated aggregates of either Tau or TAR DNA-binding protein 43 (TDP-43) [41]. This group of pathological proteinopathies causing FTD are classified under the frontotemporal lobar degeneration (FTLD) umbrella. Most cases are sporadic, however around 20% are caused by an autosomal-dominant genetic mutation including hexanucleotide repeat expansions near the chromosome 9 open reading frame gene-C9orf72, progranulin-GRN, and microtubule-associated tau protein-MAPT, as the most common causative genes [41].

FTLD pathology cause clinical bvFTD symptoms through their predominant localization in frontal and anterior temporal brain regions [41]. Clinically this is reflected by progressive cortical atrophy, which is a crucial biomarker for the diagnosis [21, 36]. While there is major overlap in atrophy patterns between sporadic and genetic bvFTD, each genetic subtype has distinctive features including antero-medial atrophy in MAPT, posterior frontal and parietal involvement in GRN and thalamic/cerebellar volume loss in C9orf72 [13]. In recent years, there has been an interest to understand how heterogeneous pathological changes could lead to similar clinical and atrophy profiles [50].

In early work based on functional magnetic resonance imaging (fMRI), it was hypothesized that atrophy in neurodegenerative diseases progresses predominantly along functional neural networks [65], with the salience network being predominantly affected in bvFTD [51, 66]. Within the salience network, the anterior insula was identified as the most likely disease epicenter [51, 65], a finding that was further supported by pathologic accumulation of tau or TDP-43 aggregates in fork cells and Von Economo neurons, which are specific to this region [32]. While the anterior insula clearly plays a significant role in the disease, other studies using data-driven methods on structural atrophy patterns revealed distinct morphological subtypes including two salience network-predominant subgroups (a frontal/temporal subtype and a frontal subtype), a semantic appraisal network-predominant group, and a subcortical-predominant group [44, 45]. This opens the possibility that there is not a single unique epicenter at the origin of all bvFTD cases.

Emerging theories emphasize that connectome architecture shapes the course and expression of multiple neurodegenerative diseases [15, 22, 30, 42, 60, 61]. Misfolding of endogenous proteins and their subsequent trans-neuronal spread has been documented in FTLD, Alzheimer’s, Parkinson’s, Huntington’s and amyotrophic lateral sclerosis (ALS) [12]. Despite differences in origin and the proteins involved in each disease, the spread of the pathology appears to reflect brain network organization at the macroscale level. Namely, anatomical connectivity is thought to support the propagation of toxic protein aggregates, such that focal pathology can spread between connected neuronal populations and infiltrate distributed networks in the brain.

Two key questions remain unanswered about the spread of pathology in bvFTD. First, the spread of pathology is likely to occur via physical white matter connections but the contribution of structural connectivity to atrophy progression has been less explored in bvFTD. Evidence for transneuronal spread of FTLD pathology is mostly based on extrapolation from functional imaging [50, 65], with some support from studies using animal models [40], autopsy data [31] and prediction of atrophy patterns [10]. Although functional connectivity reflects the underlying structural connectivity patterns and is sometimes used as a proxy for structural connectivity if no such data is available, the two modalities capture fundamentally different features of brain network organization and are only moderately correlated with each other [54]. Second, the role of local vulnerability is poorly understood. Most connectome-based models of neurodegeneration assume all neuronal populations (network nodes) have the same vulnerability to the disease, eschewing the possibility that regional differences in molecular and cellular make-up render some nodes more or less vulnerable. In particular, recent reports in other neurodegenerative diseases suggest that regional differences in gene expression may confer vulnerability, effectively guiding the pathological process through the network [23, 43, 64]. Altogether, we hypothesize that brain network architecture, in concert with local vulnerability conferred by expression of specific genes, shapes the spatial distribution of atrophy patterns in brain disorders, including FTD [51, 63, 64].

In the present report we test a structural network-based atrophy propagation model in bvFTD across sporadic and genetic variants. Specifically, we test the hypothesis that atrophy patterns in bvFTD reflect the underlying network organization and local transcriptomic vulnerability. We first estimate cortical atrophy patterns as regional changes in tissue deformation in bvFTD patients. We then use structural and functional connectivity networks derived from an independent sample of healthy individuals, to investigate whether regions that are connected with each other display similar atrophy patterns. Finally, we identify potential disease epicenters using a data-driven approach as well as a simulation-based approach that models the spread of atrophy across the brain network. We further explore the potential contribution of FTD-related genes to the propagation of atrophy.

## Results

### Demographics

Table. 1 compares demographic and clinical variables between bvFTD and CNCs across the two research databases. Subjects with bvFTDs were on average older than CNCs in the GENFI2 cohort, but not in FTLDNI. As expected, significantly lower MMSE and higher FTLD-CDR average scores were observed in symptomatic subjects compared to healthy controls.

**TABLE 1.**
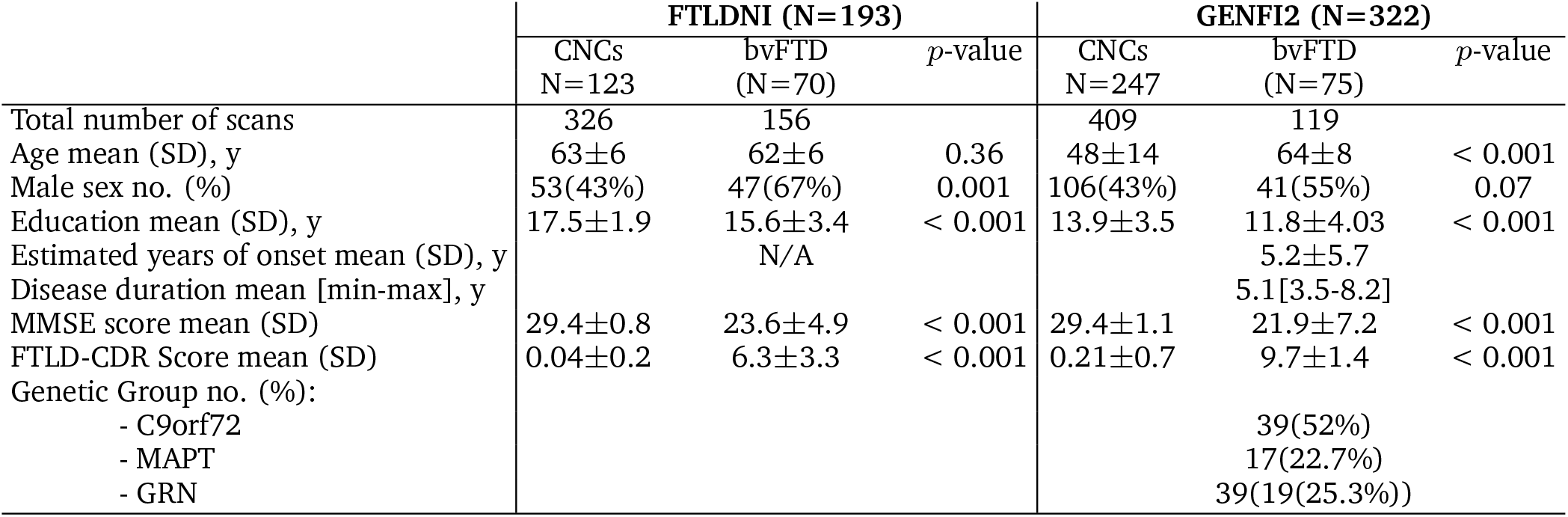
Demographic and clinical characteristics of the FTLDNI and GENFI2 samples. CNCs in GENFI2 cohort correspond to non-carrier first degree relative of a family member with a documented genetic mutation related to FTD. Genetic groups listed for CNCs in the GENFI2 cohort refer to mutation present in the family of these non-carrier subjects. Values are expressed as mean ± standard deviation, median [interquartile range]. Data available is specified for each clinical variable as N, whereas N/A indicates data not available from the original databases. (FTLDNI: frontotemporal lobar degeneration neuroimaging initiative; GENFI: genetic frontotemporal dementia initiative; bvFTD: behavioral-variant frontotemporal dementia; CNCs: cognitively normal controls; MMSE: Mini Mental State Examination. FTLD-CDR: Frontotemporal lobar degeneration clinical dementia rating.)

### Distribution of atrophy and resting state networks and cytoarchitectonic classes

We used a linear mixed effects model to obtain a group-level, bvFTD-related atrophy map, controlling for age, sex and acquisition site. The voxel-level and parcellated atrophy maps are depicted in Fig. S1a,b. In order to assess whether distributed atrophy patterns are more pronounced in specific brain systems, we used two brain system definitions (Fig. 1): (1) intrinsic functional networks defined by Yeo and colleagues [62]; (2) a cytoarchitectonic classification of human cortex based on the classic von Economo atlas [49, 56, 58, 59]. Nodes were first stratified according to their network assignments based on the Yeo networks and von Economo classes. We then calculated the mean atrophy values for each intrinsic network (Fig. 1, left) and cytoarchitectonic class (Fig. 1, right) for FTLDNI (Fig. 1a) and GENFI (Fig. 1b) datasets, separately. To assess the statistical significance of network atrophy values, we compared the empirical values to a distribution of means calculated from a set of spatial autocorrelation-preserving null models (i.e., “spin tests”[1, 35]; see *Methods* section for more details on null model). Specifically, network labels were randomly rotated while preserving the spatial autocorrelation and the mean network atrophy values were calculated for each rotation (10,000 repetitions; two-tailed test).

**Figure 1.**
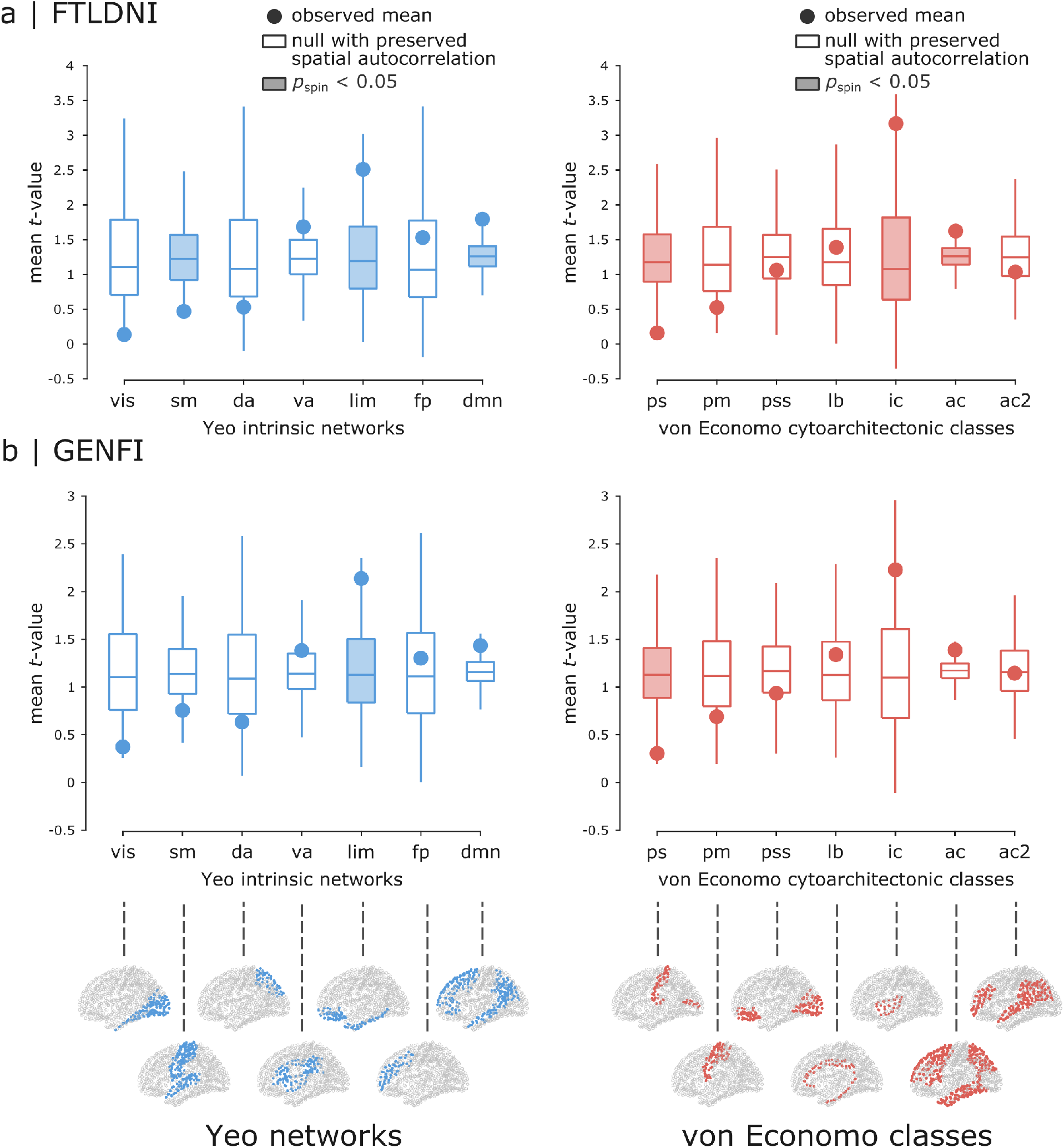
Atrophy patterns in intrinsic networks and cytoarchitectonic classes. Mean network atrophy (i.e., t-value) was calculated for Yeo intrinsic functional networks [62] (left column) and von Economo cytoarchitectonic classes [49, 56, 58, 59] (right column). Higher t-values correspond to greater atrophy. The observed mean atrophy values are shown by filled circles for each intrinsic network and cytoarchitectonic class. Network labels are then randomly permuted using 10,000 rotations from spin tests, preserving the spatial autocorrelation in the data. The null distributions of means from spin tests are depicted using box plots for intrinsic networks and cytoarchitectonic classes for both (a) FTLDNI and (b) GENFI datasets (10,000 repetitions; two-tailed test). The bottom row displays the location of intrinsic networks (left) and cytoarchitectonic classes (right) on the cortex. List of Yeo networks: visual (vis), somatomotor (sm), dorsal attention (da), ventral attention (va), limbic (lim), frontoparietal (fp), default mode (dmn). List of von Economo classes: primary sensory cortex (ps), primary motor cortex (pm), primary/secondary sensory cortex (pss), limbic (lb), insular cortex (ic), association cortex (ac, ac2).

The observed mean network atrophy and the corresponding null distribution of means are depicted for each intrinsic network and cytoarchitectonic class in Fig. 1. The anatomical distributions of intrinsic networks and cytoarchitectonic classes are depicted in Fig. 1 (bottom row). Note the difference in the definition of “limbic” system between the intrinsic networks and cytoarchitectonic classes. The intrinsic limbic network mainly consists of the temporal poles and orbitofrontal cortex, whereas the cytoarchitectonic limbic class mainly includes the cingulum. In terms of intrinsic networks, limbic and default mode intrinsic networks were the most affected (i.e., higher than expected atrophy) with relative preservation of somatomotor and visual intrinsic networks (i.e., lower than expected atrophy). In terms of cytoarchitectonic classes, the insular and association cytoarchitectonic classes displayed greater atrophy compared to nulls, with lower atrophy in primary sensory cytoarchitectonic classes. While there are marginal variations in statistical significance of the findings, the overall trend of network atrophy patterns is consistent across the two datasets.

### Relationship between atrophy maps and connectivity

We next investigated whether atrophy patterns in bvFTD are conditioned by network organization, such that connected regions display similar atrophy patterns. Specifically, we assessed whether the connectivity profile of a node can predict the atrophy of its neighbors by investigating the relationship between node and neighbor atrophy values (Fig. 2a). Structural and functional connectivity (SC and FC) networks (Fig. S1c), derived from an independent sample of 70 healthy participants [26], were used to estimate mean neighbor atrophy value for each region. The relationship between node and neighbor atrophy was then examined by correlating the mean neighbor atrophy with nodal atrophy (Fig. 2c,d). Regional atrophy was significantly correlated with the mean atrophy of its connected neighbors in both datasets. Fig. 2c (left panel) shows the results for FTLDNI dataset (high resolution parcellation: *r* = 0.69, *p*_spin_ = 0.0001 and *r* = 0.65, *p*_spin_ = 0.0001, for SC- and FC-defined neighbors respectively) and Fig. 2d (left panel) shows the results for GENFI dataset (high resolution parcellation: *r* = 0.61, *p*_spin_ = 0.001 and *r* = 0.54, *p*_spin_ = 0.0006, for SC- and FC-defined neighbors respectively). To assess whether the relationship between node and neighbor atrophy is specifically driven by network topology rather than spatial autocorrelation, we used a spatial autcorrelation-preserving null model to construct a null distribution of node-neighbor correlations [1]. Fig. 2c,d (middle panel) displays the observed correlation between node and neighbor atrophy along with the corresponding null distribution of correlations for both datasets. We also repeated all analyses at a lower parcellation resolution to ensure that the findings are robust to how network nodes are defined. The relationship between node and neighbor atrophy was consistent across resolutions and significantly greater in empirical networks compared to null networks in both datasets (Fig. 2c,d; *p*_spin_ < 0.05, two-tailed tests). The results were consistent when binarized structural connectivity network was used to defined SC-defined neighbors (Fig. S2).

**Figure 2.**
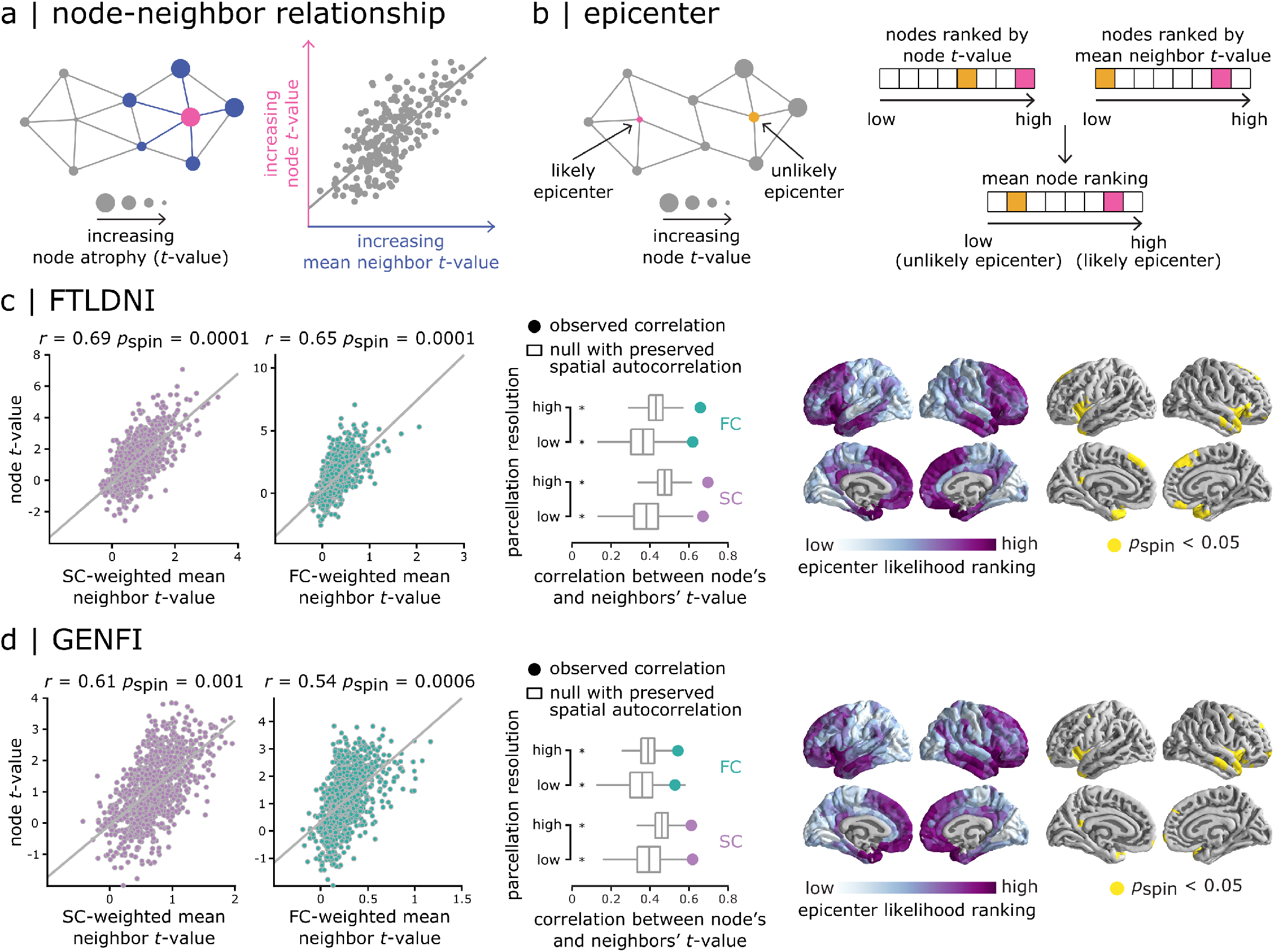
Network-dependent atrophy. (a) Atrophy of a node, estimated by t-values, was correlated with the mean atrophy of its connected neighbors to examine whether the distributed atrophy patterns in bvFTD reflect the underlying network organization. (b) If atrophy of a node is related to the atrophy of its connected neighbors (panel a), a node with high atrophy whose neighbors are also highly atrophied would be more likely to be a potential disease epicenter, compared to a high atrophy node with healthy neighbors. To quantify the epicenter likelihood across the cortex, the nodes were first ranked based on their atrophy values and their neighbors’ atrophy values. Epicenter likelihood ranking of each node was then defined as its mean ranking in the two lists. (c, d) Left panel: Node atrophy value was correlated with the mean atrophy value of its structurally- and functionally-defined neighbors (SC and FC) for FTLDNI (panel c) and GENFI (panel d) datasets. Scatter plots show the correlation for high parcellation resolution. Middle panel: The observed correlation values (depicted by filled circles) were compared to a set of correlations obtained from 10,000 spin tests (depicted by box plots). Asterisks denote statistical significance (*p*_spin_ < 0.05, two-tailed). The association between node and neighbor atrophy was consistent across resolutions and significantly greater in empirical networks compared to null networks in both datasets. Right panel: Epicenter likelihood rankings are depicted across the cortex. The most likely epicenters with high significant rankings are regions that are mainly located at the bilateral anterior insular cortex and temporal lobes (10,000 spin tests).

### Data-driven epicenters analysis

Given that the distribution of atrophy patterns reflects structural and functional network organization, we next investigated whether there are brain regions that may act as potential epicenters for bvFTD. We define an epicenter as a high atrophy node that is connected to high atrophy neighbors (Fig. 2b). Nodes were ranked based on their atrophy and their neighbors’ mean atrophy values. Epicenter likelihood ranking was then estimated as the mean node ranking across the two lists. Fig. 2c,d (rightmost panel) shows the epicenter likelihood rankings on the cortex for FTLDNI (Fig. 2c) and GENFI (Fig. 2d) datasets, where the highly ranked regions are associated with insular cortex, ventromedial cortex and antero-medial temporal areas. Empirical epicenter likelihood rankings were then compared with rankings estimated from spatial autocorrelation-preserving null models (10,000 spin tests [1]). Several regions were identified as potential epicenters including the anterior insular cortex bilaterally, but also areas in the anterior temporal poles, in addition to ventromedial and dorsomedial areas. The results were consistent when binarized structural connectivity network was used to defined SC-defined neighbors (Fig. S2).

### Dynamic spreading model

We next used an S.I.R model to explore how the brain’s structural connectivity shapes the progressive spread of FTLD changes. This model has been previously used to study Parkinson’s disease-related atrophy [64] and works by simulating the misfolding of normal proteins in the cortex and their trans-neuronal spread through the structural connections between brain regions. The accumulation of misfolded proteins, acting as pathogenic agents, leads to the atrophy of the afflicted regions (Fig. 3a). Epicenters are defined as those regions in which misfolded proteins are introduced, allowing us to test which is the most likely epicenter for the observed empirical patterns. As misfolded agents spread through the network, we measure the Pearson correlation between the simulated and empirical (FTLDNI) patterns of atrophy (Fig. 3b; left panel). A region’s maximal correlation (*r*_max_) is defined as the largest correlation value observed across all values of t. The three nodes with the largest *r*_max_ are located in the insular, superior-frontal and lateral orbito-frontal cortex.

**Figure 3.**
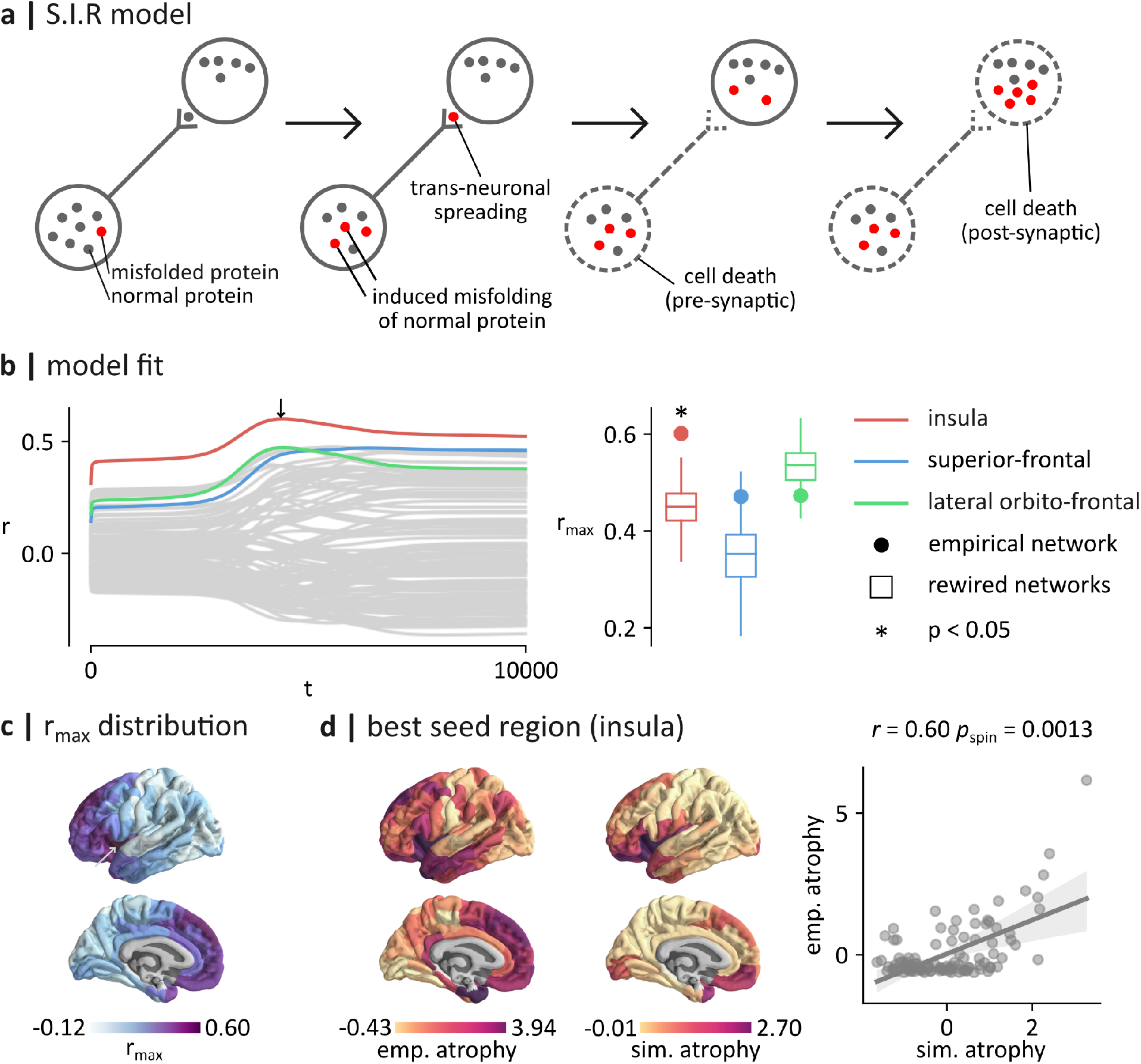
Agent-based modeling. (a) The S.I.R model simulates the spread of pathogenic agents (misfolded proteins) in the brain. Proteins propagate via the structural connections between brain regions and induce atrophy, both pre- and post-synaptically. (b) Left panel: the spreading process was initiated in every brain region and the correlation between the simulated and empirical patterns of atrophy was computed. The three largest correlations were obtained by seeding regions of the insula (*r*_max_ = 0.601; red), the superior-frontal cortex (*r*_max_ = 0.473; blue) and lateral orbito-frontal cortex (*r*_max_ = 0.471; green). The correlations for other brain regions are shown in gray. Right panel: to control for the potential effect of a brain region’s spatial embedding, *r*_max_ values were compared to *r*_max_ correlations obtained using rewired networks that preserve the wiring-cost of the empirical structural network. Asterisks denote statistical significance (*p*< 0.05, two-tailed). The *r*_max_ computed by seeding the insula of the empirical network (*r*_max_ = 0.60) was significantly larger than the *r*_max_ computed by seeding the insula of the rewired networks (*p* < 0.002). (c) The largest fit (*r*_max_) obtained by seeding each brain region is shown on the surface of the brain. Larger values of *r*_max_ were generally obtained by seeding insular and prefrontal regions. (d) Left panel: empirical pattern of atrophy (FTLDNI). Middle panel: simulated pattern of atrophy producing the maximal fit. This pattern of atrophy was obtained with the insula as the seed, and at t=4410 (see the arrow in panel b). Right panel: scatterplot of the relationship between standardized empirical and simulated patterns of atrophy (*r* = 0.60, *p*_spin_ = 0.0013).

An important factor that can influence the probability that a brain region is identified as the epicenter of an atrophy pattern is its spatial location in the brain. To isolate the role of structural connectivity, we compared these *r*_max_ scores to those obtained by simulating the spread of misfolded proteins in rewired networks that preserve the density, degree sequence and wiring cost of the empirical structural network (Fig. 3b; right panel). We find that the fit obtained by initiating the spread in the insular region of the empirical network is significantly larger than the fit obtained in the rewired networks (*r* = 0.601, *p* < 0.002). In other words, the fit observed by seeding the insula is significantly larger than what would have been expected from its degree and spatial position alone and can be attributed to its embedding in the global topology of the network. This result suggests that the topology of the structural connectome plays a significant role in shaping patterns of simulated atrophy that have a high correspondence with the empirical atrophy.

More generally, by looking at the topographic distribution of *r*_max_ scores, we find that the brain regions that show the largest fits are located in the insular, medial prefrontal and anterior temporal cortices (Fig. 3c). These results are in accordance with our finding that these regions have large epicenter likelihood rankings. Fig. 3d shows the empirical pattern of atrophy for the FTLDNI dataset. This pattern is compared to the simulated pattern of atrophy producing the maximal fit. This largest fit was obtained by seeding the insula and was measured at *t* = 4410. We find a significant relationship between the two distributions (*r* = 0.60, *p*_spin_ = 0.0013). Results are presented for the FTLDNI dataset, but similar results are found in the GENFI dataset (Fig. S3).

### Contribution of gene expression to network spreading

Given the contribution of genetic variants to bvFTD [25], we next assessed whether the incorporation of gene expression information into the S.I.R model can enhance the fits. We used regional microarray expression data from the Allan Human Brain Atlas [28] to generate vectors of gene expression for four genes that have been previously associated with bvFTD: MAPT, GRN, C9orf72 and TARDBP [41]. Fig. 4a shows the topographic distributions of these genes. We used this genetic information to set regional heterogeneity for the clearance and synthesis of proteins in the model. We used the insula as the seed region of the spreading process as it is the region that showed the largest fit to the empirical data.

**Figure 4.**
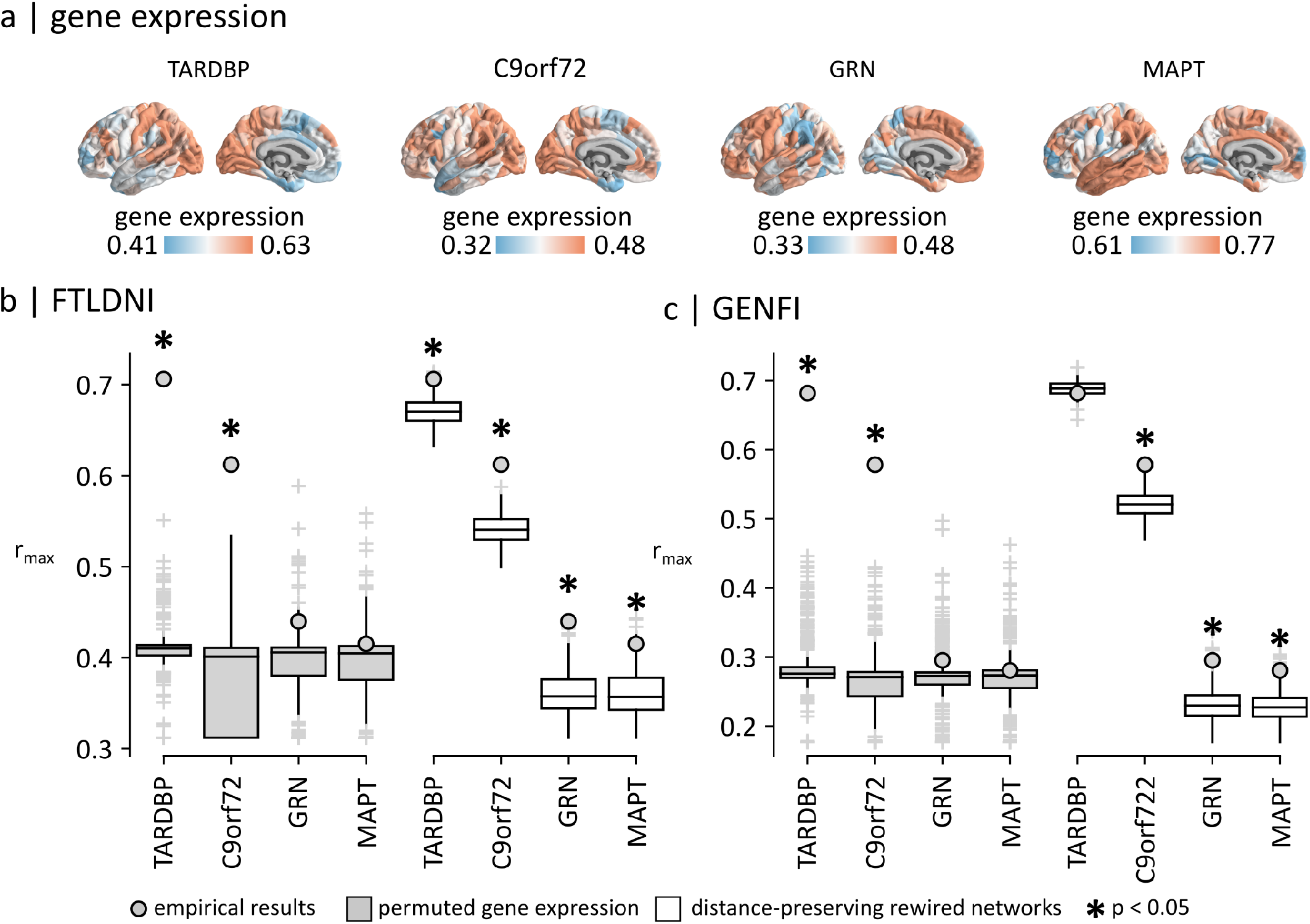
Contribution of gene expression. (a) Vectors of regional gene expression were generated for four genes that have been associated with bvFTD: TARDBP, C9orf72, GRN and MAPT. These vectors of gene expression were incorporated into the S.I.R model. The correlations between empirical atrophy and simulated atrophy, with the insula selected as the seed of the simulated spreading process, were then computed for the FTLDNI dataset (b) and for the GENFI dataset (c). The maximal correlations scores (*r*_max_) obtained for each gene were compared to the maximal correlation scores (*r*_max_) obtained with permuted gene expression vectors (grey boxplots). Asterisks denote statistical significance (*p* < 0.05, two-tailed). For both datasets, we find that the *r*_max_ scores obtained by incorporating information about the expression of C9orf72 and TARDBP are significantly larger than those obtained with permuted gene expression vectors (*p* < 0.0001). The maximal correlations are also compared to the maximal correlation scores obtained in null networks (white box plots).

For both FTLDNI (Fig. 4b) and GENFI (Fig. 4c) datasets, we measured the *r*_max_ scores obtained by incorporating regional expression for each of the four genes. With MAPT, GRN, C9orf72 and TARDBP, we obtain correlation scores of *r*_max_ = 0.42, *r*_max_ = 0.44, *r*_max_ = 0.61 and *r*_max_ = 0.71 for the FTLDNI dataset, and *r*_max_ = 0.28, *r*_max_ = 0.30, *r*_max_ = 0.58 and *r*_max_ = 0.68 for the GENFI dataset. We find that adding regional heterogeneity for synthesis and clearance using expression of C9orf72 and TARDBP increased model fit while the incorporation of information regarding the regional expression of GRN and MAPT decreased model fit. To investigate the significance of the findings, we randomly permuted the vectors of gene expression 10,000 times to generate null distributions of *r*_max_ scores that we compared to the empirical results. We find that the scores obtained with C9orf72 and TARDBP are significantly larger than those obtained with permuted gene expression vectors (*p*< 0.0001). These results suggest that C9orf72 and TARDBP may play a significant role in driving the spatial patterning of the empirical atrophy.

To investigate the relationship between gene expression and the brain’s structural connectivity, we compared the fits to those obtained using rewired networks preserving the wiring-cost of the empirical network. For C9orf72, we find that the fits obtained using the empirical networks were significantly larger than the fits obtained using rewired null networks, for both FTLDNI (*p*< 0.002) and GENFI (*p*< 0.002). This result suggests that for both sporadic bvFTD and genetic bvFTD, the topology of the structural connectome has a positive influence on the increase in model fit observed when incorporating C9orf72 into our model. For TARDBP, we find that the fits obtained using the empirical networks were significantly larger than the fits obtained using rewired null networks for FTLDNI (*p* = 0.014), but not for GENFI (*p* = 0.508). This result suggests that the topology of the structural connectome appears to have a positive influence on the increase in model fit observed when trying to fit our model to empirical atrophy patterns associated to sporadic bvFTD, but not when trying to fit our model to empirical atrophy patterns associated to genetic bvFTD.

## Discussion

The present report provides a comprehensive and statistically robust model supporting the theory of network-based atrophy in bvFTD, both in sporadic and genetic forms. Our findings are consistent across two datasets and the genetic/sporadic heterogeneity. Namely, for both sporadic and genetic variants, there is a strong correlation between node deformation and the mean of neighbor deformation defined by both structural and functional connectivity, supporting the theory that atrophy progresses through network-based connections. Similar findings were observed at small (219) and large (1000) cortical parcellation resolutions. Data-driven epicenter mapping identified the bilateral anterior insular cortex, as well as ventromedial cortex and antero-temporal areas as potential epicenters. The involvement of the antero-medial areas as epicenters ties into previous research showing that data-driven atrophy subtypes include a “semantic appraisal network” predominant group [27, 44]. The genetic bvFTD cohort showed a very similar profile of most likely epicenters, with the addition of some dorsal frontal areas. The role of these regions as epicenters was further supported by the agent-based spreading model.

The localization of cortical atrophy was most significant in the limbic resting state network, and less present in the visuospatial network (expectedly given its posterior localization). There was significant atrophy in the default mode network (DMN) in genetic FTD, with a positive trend in sporadic FTD. Of note, the salience network which has been previously identified as being predominantly involved in bvFTD [51, 66] did not show statistically significant atrophy. However, when looking at von Economo cytoarchitectonic classes, the insular cortex was the most affected, with relative sparing of the primary sensory neurons. This suggests that the insular cortex plays a central role in the disease, but not necessarily by spreading through the entire ventral attention network including its most posterior regions. In addition, while there have been some reports of opposite connectivity pattern of changes in the salience versus the DMN in bvFTD and AD [66], our results rather suggest that there is significant involvement of DMN regions in bvFTD.

Finally, although exploratory, using a simulation-based approach and gene expression profile data from the Allen Human Brain Atlas, we identified that the C9orf72 and TARDPB gene expression could play a role in the propagation of atrophy in sporadic bvFTD. Indeed, factoring an impact on clearance and synthesis of both genes related to TDP-43 improved the fit between the modeling spreading models and the actual atrophy maps based on DBM. While we cannot exclude that some subjects in the FTLDNI had an unidentified C9orf72 mutation, the involvement of TARDPB is of interest given that that mutations in this gene constitute only a very small fraction of genetic FTD. Results suggest that the activity of this gene could play a role in sporadic bvFTD, which could be of interest for future therapeutic avenues. In future studies, it would be of interest to model the role of genes in the spread of atrophy within each genetic subgroup separately, including all clinical phenotypes.

This study has several strengths including the replication across two separate samples of genetic and sporadic bvFTD. DBM is a robust method to capture atrophic changes in patients with neurodegenerative dementia. The present findings were replicated in two separate samples of genetic and sporadic bvFTD and were validated using a range of methodological choices. We also confirmed that the findings are independent from potential confounding factors such as spatial distance and parcellation resolution. However, there are several methodological considerations that need to be taken into account when interpreting the findings. First, there are no available molecular techniques to directly measure FTLD changes in vivo. To overcome this limitation, we opted to use DBM to estimate atrophy in bvFTD patients since it is a robust method to capture local changes in brain tissue volume. Second, diffusion spectrum imaging and streamline tractography were used to estimate structural connectivity networks. Although this approach is widely applied to reconstruct human structural connectome using non-invasive neuroimaging data, it is prone to false positives and negatives in defining white matter connections [29, 55]. Third, we identified potential disease epicenters using cross sectional data and undirected networks, therefore it is not possible to precisely assess the propagation process of pathology across the connectome architecture. Modeling disease progression and spread of atrophy across brain networks over time remains an exciting open question that could potentially be addressed by investigating a longitudinal sample of FTD patients using the methods presented here. Fourth, the two datasets included in this study have different demographics and clinical variables (e.g., age range, sex, education, disease severity, etc.). However, the results were mainly consistent across the two datasets, confirming that the findings are robust against those differences. Finally, the structural and functional connectivity networks used in this study are measured in a sample of young healthy population rather than FTD patients. We used these networks as a healthy underlying architecture that may allow pathogen transmission between the connected regions. However, it is possible that the connectome architecture itself would also be compromised during the course of the disease progression, rerouting or restricting the spread of pathology. This could be further investigated in future studies using simultaneous measures of regional atrophy and changes in structural and functional connectivity in a longitudinal sample of FTD patients.

### Summary

Altogether, structural and functional connectivity networks and rigorous statistical analyses that account for spatial autocorrelation and network embedding are used in the present study to demonstrate that bvFTD-related neurodegeneration is conditioned by connectome architecture, accounting for 30-40% of variance in atrophy, as well as local transcriptomic vulnerability. FTD-related atrophy appears to be particularly targeting regions associated with the anterior insular cortex, but it is likely that there are multiple potential epicenters leading to bvFTD clinical phenotypes. The similarity between genetic and sporadic forms of bvFTD suggests that multiple pathological changes are constrained by the network architecture in the spread of atrophy, explaining why many different pathological and genetic entity lead to the same clinical syndrome. Although exploratory, our results suggest that TARDPB gene expression could have a significant contribution to disease progression, particularly in sporadic bvFTD.

## Methods

### Participants

We retrieved data from subjects with bvFTD and cognitively normal controls (CNCs) from the Frontotemporal Lobar Degeneration Neuroimaging Initiative (FTLDNI) database that had T1-weighted (T1w) MRI scans matching with each clinical visit (http://4rtni-ftldni.ini.usc.edu/). The inclusion criteria for bvFTD patients were a diagnosis of possible or probable bvFTD according to the FTD consortium criteria [46], resulting in 70 patients with bvFTD (mostly sporadic) and 123 CNCs available for analyses. Several patients had more than one scan therefore there was a total of 156 scans in the bvFTD group and the 326 in the CNC group. We also accessed data from the third data freeze (12/2017) of the Genetic Frontotemporal Dementia Initiative 2 (GENFI2 -http://genfi.org.uk/), which includes 23 centers in the UK, Europe and Canada [48]. GENFI2 participants include known symptomatic carriers of a pathogenic mutation in C9orf72, GRN or MAPT and their first-degree relatives who are at risk of carrying a mutation, but who did not show any symptoms (i.e., at-risk subjects). Healthy first-degree relatives who were found to be non-carriers of a mutation are considered as CNCs. Since the aim of the present study was to study network propagation of atrophy in the bvFTD clinical phenotype, presymptomatic carriers and symptomatic carriers whose clinical diagnosis was other than bvFTD were excluded. This GENFI2 cohort included 75 patients with bvFTD and 247 CNCs. Demographic and clinical characteristics of those two cohorts are described in Table. 1. Two-sample t-tests were conducted to examine demographic and clinical variables at baseline. Categorical variables were analyzed using chi-square analyses. Results are expressed as mean ± standard deviation and median [interquartile range] as appropriate.

### MRI acquisition and processing

For the FTLDNI cohort, 3.0T MRIs were acquired at three sites (T1w MPRAGE, TR=2 ms, TE=3 ms, IT=900 ms, flip angle 9^*o*^, matrix 256×240, slice thickness 1mm, voxel size 1mm^3^). For the GENFI2 sample, volumetric T1w MPRAGE MRI was obtained at multiple centers using the GENFI imaging protocol on either Siemens Trio 3T, SiemensSkyra3T, Siemens1.5T, Phillips3T, General Electric (GE) 1.5T or GE 3T scanners. Scan protocols were designed at the outset of the study to ensure adequate matching between the scanners and image quality control.

All T1w scans were pre-processed through our longitudinal pipeline that includes image denoising, intensity non-uniformity correction and image intensity normalization into range (0-100) using histogram matching [5, 14, 16, 53]. The image processing tools used in this study were designed to process data from multisite studies to handle biases due to multi-site scanning and they have been successfully applied to a number of multi-site projects [9, 17, 18, 63]. Each native T1w volume from each time point was linearly registered first to the subject-specific template which was then registered to the ICBM152 template. All images were then non-linearly registered to the ICBM152 template using ANTs diffeomorphic registration pipeline [6]. The images were visually assessed by two experienced raters (MD and ALM) to exclude cases with significant imaging artifacts (e.g., motion, incomplete field of view) or inaccurate linear/nonlinear registrations. This visual quality control was completed blind to the diagnosis. Out of 1724 scans, only 43 (2.5%, 36 scans in GENFI2, and 7 in FTLDNI) were rejected. This resulted in a total of 515 subjects that were included to perform cross-sectional morphometric analyses.

### Deformation-based morphometry (DBM) analyses

DBM [3, 4] analysis was performed using Montreal Neurological Institute (MNI) MINC tools [33]. The local deformations, obtained from the non-linear transformations mapping the MNI-ICBM152-2009c template to the subject’s MRI, encode the local tissue volume difference between the MNI average template and subject’s brain. The determinant of the Jacobian of the deformation field is measured at each voxel. Determinant values larger than 1.0 indicate that the local volume in the subject is larger than the average template (e.g., ventricular or sulci enlargement in the case of FTD). Determinant values smaller than one indicate that the local volume in the subject is smaller than the template. The latter is often interpreted as tissue atrophy despite the use of only cross-sectional data. DBM was used to assess voxel-wise cross-sectional group related volumetric differences. To obtain a voxel-wise map reflecting the patterns of difference between bvFTD and CNCs, the following mixed effects model was applied on a voxel-by-voxel basis, separately for each dataset:

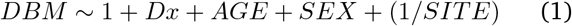

The mixed effects model included age as a continuous fixed variable and diagnosis (*Dx*) and sex as fixed categorical variables. Site was included as a categorical random variable. The variable of interest was diagnosis, reflecting the brain regions that were significantly different between bvFTD and CNCs, controlling for age and sex. Statistical t-maps were extracted from the model and used for the rest of the analyses throughout the manuscript. Finally, the t-statistics were multiplied by -1 such that higher positive values correspond to higher atrophy and negative values correspond to volume expansion.

### Anatomical parcellation

Statistical t-maps obtained through DBM analysis and mixed effects models were parcellated into 219 and 1000 approximately equally sized cortical regions or parcels using the Cammoun atlas [11], a multiresolution extension of the anatomical Desikan-Killiany atlas [20]. We refer to 219 and 1000 parcellation resolutions as low and high parcellation resolutions, respectively. The parcelwise t-statistics (i.e., atrophy) were estimated as the mean t-statistic of all the voxels that were assigned to that parcel according to the atlas. We repeated all the analyses at both parcellation resolutions to ensure that results are replicable across multiple spatial scales.

### Structural and functional network reconstruction

Connection patterns from healthy individuals are used to represent the architecture of brain networks for the distributed atrophy patterns that are observed in bvFTD patients. Structural and functional connectivity data of 70 healthy individuals (mean age 28.8 ± 9.1 years) were obtained from a publicly available dataset [26]. Details about data acquisition parameters and pre-processing analysis are available in [26]. Briefly, the participants were scanned in a 3T MRI scanner (Trio, Siemens Medical, Germany) using a 32-channel head-coil. The session protocol included: (1) a magnetization-prepared rapid acquisition gradient echo (MPRAGE) sequence sensitive to white/gray matter contrast (1-mm in-plane resolution, 1.2-mm slice thickness); (2) a DSI (diffusion spectrum imaging) sequence (128 diffusion-weighted volumes and a single *b*_0_ volume, maximum *b*-value 8,000 s/mm^2^, 2.2×2.2×3.0 mm voxel size); and (3) a gradient echo EPI sequence sensitive to BOLD (blood-oxygen-level-dependent) contrast (3.3-mm in-plane resolution and slice thickness with a 0.3-mm gap, TR 1,920 ms, resulting in 280 images per participant). Diffusion spectrum imaging data and deterministic streamline tractography were used to construct structural connectivity networks for each healthy individual. Each pair-wise structural connection was weighted by the log-transform of the fiber density. Individual structural connectivity networks were parcellated into the low and high parcellation resolutions using the Cammoun atlas described before. Resting-state functional MRI data collected in the same healthy individuals (with eyes open) were used to construct functional connectivity networks. The preprocessed resting-state functional MRI time series were also parcellated using both the low and high resolution versions of the Cammoun atlas and were correlated to estimate functional connectivity between pairs of brain regions using Pearson correlation coefficients. Finally, a consensus group-average structural connectivity preserving the edge length distributions in individual networks was constructed [8, 37, 38] and a group-average functional connectivity was estimated as the mean pairwise connectivity across individuals.

### Network atrophy

Group-average structural and functional connectivity networks were used to estimate average atrophy values of neighbors of each brain region [52]. Briefly, neighbors of a given brain region were defined as regions that are connected to it with a structural connection for both structurally- and functionally-defined neighbors. The structurally-connected neighbor atrophy value of each brain region was then estimated as the average weighted atrophy values of all the neighbors of that region:

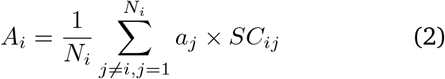

where *A*_*i*_ is the average neighbor atrophy value of brain region or node *i, a*_*j*_ is atrophy of *j*-th neighbor of node *i, SC*_*ij*_ is the strength of structural connection between nodes *i* and *j*, and *N*_*i*_ is the total number of neighbors that are connected to node *i* with a structural connection (i.e., node degree). Normalization by term *N*_*i*_ ensures that the estimated neighbor atrophy value is independent from the node degree. The neighbor atrophy estimation excludes self-connections (*j* ≠ *i*). The functionally-connected neighbor atrophy values were estimated using the same equation as above, with the exception that regional atrophy values were weighted by the strength of functional connections between nodes *I* and *j* (*FC*_*ij*_):

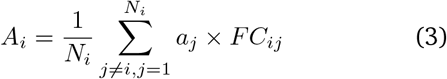

For both structurally- and functionally-defined neighbor atrophy estimates, neighbors were defined as nodes that were structurally connected to the node under consideration. Altogether, a single neighbor atrophy value was estimated for each region. We used Pearson correlation coefficients to assess the relationship between node atrophy and mean neighbor atrophy for structurally- and functionally-defined neighbors, separately (Fig. 2a).

### Data-driven epicenter analysis

To identify potential disease epicenters, we hypothesized that an epicenter would be a node with high atrophy that is also connected to highly atrophied neighbors, compared to a high atrophy node with healthy neighbors, or a healthy node with atrophied neighbors. Using a data-driven approach [52], we first ranked the nodes based on their estimated regional atrophy values. We also ranked the nodes based on the average atrophy values of their neighbors in a separate list. We then calculated the average ranking of each node in the two lists and identified nodes that were highly ranked in both lists (i.e., nodes with both high local and neighborhood average rankings) as the potential epicenters (Fig. 2b).

### Agent-based spreading model

#### Simulation-based epicenter analysis

To investigate the transneuronal spread hypothesis, we simulated the spread of misfolded proteins on the left hemisphere of the low-resolution weighted consensus structural connectivity network (111 regions) using a Susceptible-Infected-Removed (S.I.R) agent-based model [64]. Briefly, the model consists of simulating the misfolding of normal proteins in the cortex and their trans-neuronal spreading through the structural connections between brain regions. The accumulation of misfolded proteins, which act as pathogenic agents, then leads to the atrophy of the afflicted regions. Importantly, this model incorporates synthesis and clearance rates, which can heterogeneously vary across brain regions. More details about the model’s main equations can be found in the *Supplemental Information*. To explore the likelihood that a brain region acts as an epicenter of this spreading process, we first used baseline clearance and synthesis rates for all regions. We simulated the spread of misfolded proteins and the resulting atrophy using, one at a time, each individual brain region as the seed of the process. For each seed region, and at each time point, we then computed the Pearson correlation between the simulated and empirical patterns of atrophy.

#### Gene expression

To investigate the potential role of gene expression in shaping the modelled patterns of atrophy, we accessed the Allen Human Brain Atlas (AHBA [28]; http://human.brain-map.org/), which provides regional microarray expression data from six post-mortem brains (1 female, ages 24-57, 42.5 ± 13.38). We used the atlas to generate vectors storing gene expression scores for each of the 111 regional parcel of the left hemisphere. More specifically, we focused on four vectors of gene expression associated with genes that have been previously linked to bvFTD, namely MAPT, GRN, C9orf72 and TARDBP. Given that subjects were selected based on their clinical phenotype (bvFTD) rather than on a specific pathological subtype or genetic mutation, we explored the potential role of the expression of all four genes for both synthesis and clearance. As such, vectors of gene expression were used to specify the synthesis and clearance rate of each brain region, such that a greater expression score entailed a greater synthesis and clearance rate. Our objective was to identify potentially new mechanistic processes underlying the spreading of atrophy, particularly in sporadic bvFTD where we do not have adequate knowledge of the contribution of genes related to the various proteinopathies.

The AHBA data was pre-processed and mapped to the parcellated brain regions using the abagen toolbox [34] (https://github.com/rmarkello/abagen). During pre-processing, we first updated the MNI coordinates of tissue samples to those generated via non-linear alignment to the ICBM152 template anatomy (https://github.com/chrisgorgo/alleninf). We also reannotated the microarray probe information for all genes using data provided by [2]. We then filtered the probes by only retaining those that have a proportion of signal to noise ratio greater than 0.5. When multiple probes indexed the expression of the same gene, we selected the one with the most consistent pattern of regional variation across donors. Samples were then assigned to individual regions in the Cammoun atlas. If a sample was not found directly within a parcel, the nearest sample, up to a 2mm-distance, was selected. If no samples were found within 2mm of the parcel, we used the sample closest to the centroid of the empty parcel across all donors. To reduce the potential for misassignment, sample-to-region matching was constrained by hemisphere and gross structural divisions (i.e., cortex, subcortex/brainstem, and cerebellum, such that e.g., a sample in the left cortex could only be assigned to an atlas parcel in the left cortex). All tissue samples not assigned to a brain region in the provided atlas were discarded. Tissue sample expression scores were then normalized across genes using a scaled robust sigmoid function [24], and were rescaled to a unit interval. Expression scores were also normalized across tissue samples using the same procedure. We then aggregated the microarray samples belonging to the same regions by computing the mean expression across samples for individual parcels, for each donor. Regional expression profiles were finally averaged across donors.

### Null models

To assess the statistical significance of the node-neighbor relationships and the epicenter analysis, we used a spatial autocorrelation preserving null model (i.e., “spin tests” [1, 35]). We first used the Connectome Mapper toolkit ([19]; https://github.com/LTS5/cmp) to generate a surface-based representation of the Cammoun atlas (both low and high resolution) on the Freesurfer fsaverage surface. We then defined the spatial coordinates of each parcel by selecting the vertex on the spherical projection of the generated fsaverage surface that was closest to the center of mass of the parcel [52, 57]. Finally, we used the resulting parcel spatial coordinates to generate null models of brain maps (e.g., atrophy maps, epicenter rankings) by randomly rotating the maps and reassigning node values with the values of closest parcels. The rotations were first applied to one hemisphere and the mirrored rotations were used for the other hemisphere. This procedure was repeated 10,000 times to generate a null distribution of brain maps with preserved spatial autocorrelation.

To ensure that the observed correlation between the empirical and simulated atrophy map from the agent-based model is explained by the topological organization of the structural connection between brain regions and not solely by the spatial embedding of brain regions, we generated surrogate networks that preserve the geometry of the structural connectome. The edges of the consensus network were first binned according to inter-regional Euclidean distance. Within each length bin, pairs of edges were then selected at random and swapped [7]. This procedure was repeated 500 times, generating a population of rewired structural networks that preserve the degree sequence of the original network and that approximately preserve the edge length distribution (i.e., wiring cost) of the empirical network.

## Acknowledgments

This research was undertaken thanks in part to funding from the Canada First Research Excellence Fund, awarded to McGill University for the Healthy Brains for Healthy Lives initiative. BM acknowledges support from the Natural Sciences and Engineering Research Council of Canada (NSERC Discovery Grant RGPIN #017-04265) and from the Canada Research Chairs Program. GS acknowledges support from the Natural Sciences and Engineering Research Council of Canada (NSERC) and the Fonds de recherche du Québec - Nature et Technologies (FRQNT).

## Supplemental Information

### The S.I.R model

The S.I.R model has been previously used to explore the spreading of pathological proteins in Parkinson disease [64]. The model is described at length in the original paper. The code of the model can be found at https://github.com/yingqiuz/SIR_simulator. This section briefly summarizes the model’s main equations.

The initial step of this model consists in determining the baseline regional density of some protein of interest in individual parcels of the network. To do so, we increment the population of normal agents (i.e., proteins) in region *i, N*_*i*_, with

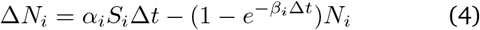

where *α*_*i*_ and *β*_*i*_ are respectively the synthesis and clearance rate of the protein in region *i. S*_*i*_ corresponds to the size (voxel count) of region *i* and Δ*t* is the time interval between two iterations, which is set to 0.02.

After the system reaches the stable point (error tolerance ϵ < 10^*-*7^), the pathogenic spread of misfolded proteins is initiated and the population of normal (*N*) and misfolded (*M*) agent is updated with

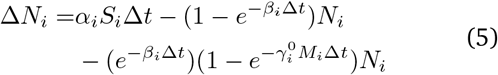

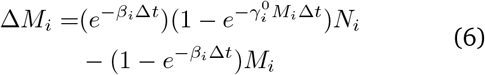

where 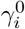 is the baseline transmission rate that measures the likelihood that a single misfolded agent can transmit the infection to other susceptible agents. This baseline transmission rate was set to 1*/S*_*i*_.

For each iteration, agents in region *i* may remain in the region or may enter one of its edges according to a multinomial distribution with probabilities

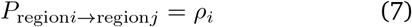

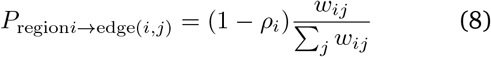

where *w*_*ij*_ is the connection strength of edge (*i, j*). The probability of remaining in the current region, *ρ*_*i*_, was set to 0.99. Analogously, the agents in edge (*i, j*) may exit the edge or remain in the same edge per unit of time with binary probabilities:

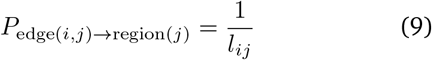

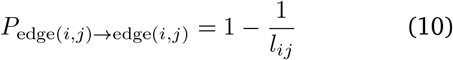

where *l*_*ij*_ is the length of edge (*i, j*).

We use *N*_*i,j*_ and *M*_*i,j*_ to denote the normal/misfolded population in edge (*i, j*). For each interval of time Δ*t*, the increments of *N*_*i*_, *M*_*i*_ in region *i* are

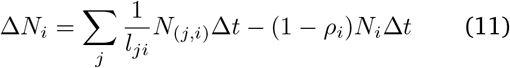

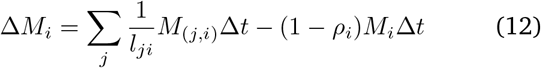

Likewise,

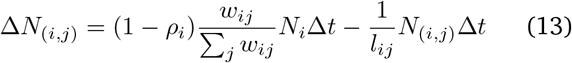

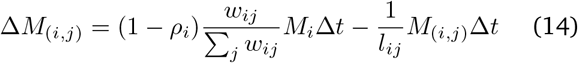

Finally, we model neuronal tissue loss (*L*) as the result of 2 processes: direct toxicity from accumulation of native misfolded proteins and deafferentation (reduction in neuronal inputs) from neuronal death in neighboring (connected) regions. The atrophy accrual at *t* within Δ*t* in region *i* is given by the sum of the two processes:

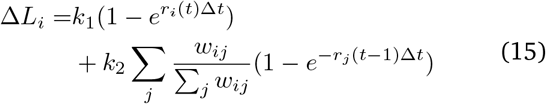

Where *r*_*i*_(*t*) is the proportion of misfolded agents in region *i* at time *t. k*_1_ and *k*_2_ are both set to 0.5 such that the 2 processes have equal importance in modelling the total atrophy growth. A list of the free parameters that have been used can be found in Table. S1.

**TABLE S1.** Parameters of the S.I.R model. Values of the free parameters of the S.I.R model used in this study are listed. For a complete list of all the parameters or notations used in the model, see [64].

| Notation | Name | Expression or value | Explanation |
| --- | --- | --- | --- |
| $\Delta t$ | time step | $\Delta t = 0.02$ | Time increments in the simulations |
| $\alpha_i$ | synthesis rate in region $i$ | See *Methods* section | Probability that a new normal agent gets synthesized in each voxel of region $i$ per unit time |
| $\beta_i$ | clearance rate in region $i$ | See *Methods* section | Probability that an existing agent (either normal or misfolded) in region $i$ gets cleared per unit time |
| $\rho_i$ | the probability of remaining in region $i$ | $\rho_i = 0.99$ for all $i$ | Agents in region $i$ have equal probability of remaining in region $i$ or exiting region $i$ per unit time |
| $k_1$ | weight of atrophy accrual due to accumulation of misfolded agents | $k_1 = k_2 = 0.5$ | The contribution of native misfolded agents to total atrophy growth |
| $k_2$ | weight of atrophy accrual due to deafferentation | $k_1 = k_2 = 0.5$ | The contribution of deafferentation to total atrophy growth |
| $w_{ij}$ | Connection strength between regions $i$ and $j$ | Log-transform of streamline density | |
| $l_{ij}$ | Connection length between regions $i$ and $j$ | Euclidean length of streamline fibers | |

**Figure S1.**
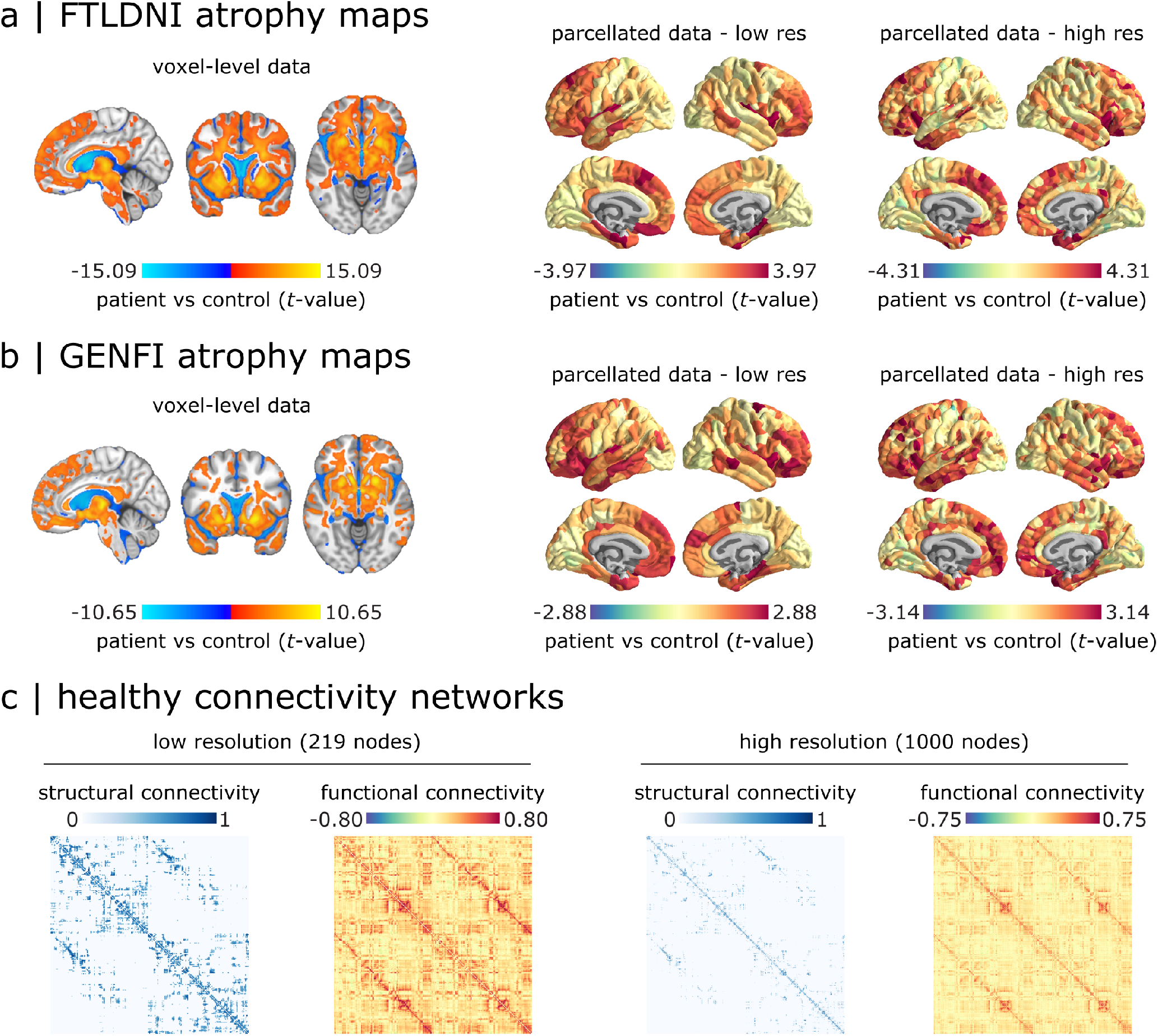
bvFTD atrophy maps and healthy connectivity networks. Voxel-level and parcellated atrophy maps are shown for (a) FTLDNI and (b) GENFI datasets. The voxel-level data are displayed on the MNI-ICBM152 template (*x* = − 5, *y* = 13, *z* = − 6). Greater t-values correspond to greater atrophy in bvFTD patients relative to healthy control subjects. The parcellated data are displayed on brain surface at 95% intervals for both low and high parcellation resolutions (219 and 1000 nodes, respectively). (c) Group-average structural and functional connectivity networks, derived from an independent sample of 70 healthy participants [26], are shown for low and high parcellation resolutions. Each pair-wise structural and functional connection correspond to the mean log-transform of the fiber density and the mean Pearson correlation coefficient between regional resting-state functional MRI time series across individuals, respectively.

**Figure S2.**
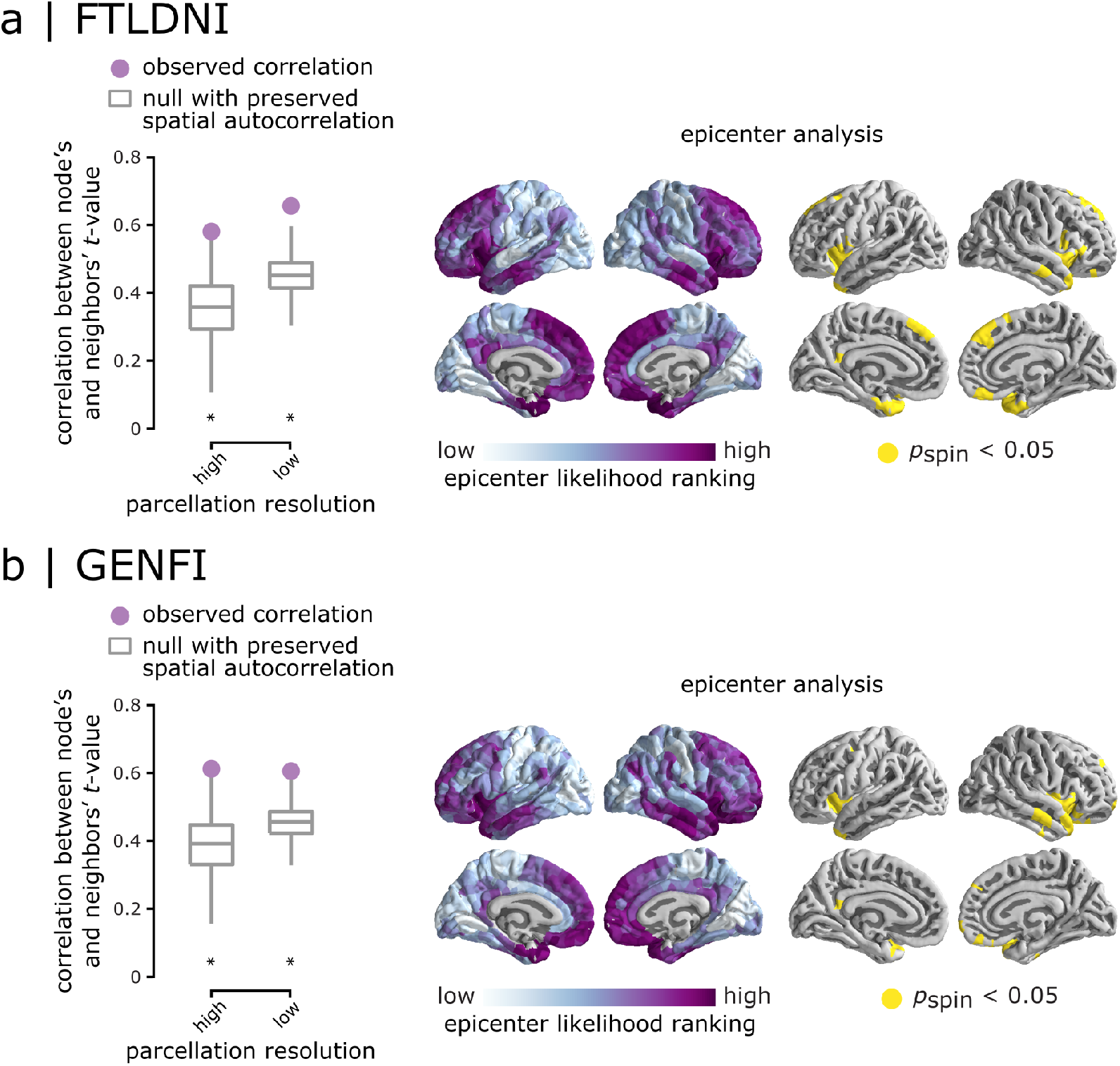
Network-dependent atrophy using binary structural connectivity. Binarized structural connectivity was used to estimate the SC-defined mean neighbor atrophy (i.e., t-value) for (a) FTLDNI and (b) GENFI datasets. More specifically, the neighbor atrophy values were not weighted by the strength of structural connectivity. Left panel: Atrophy of a node was correlated with the mean atrophy of its connected neighbors. The observed correlation values (depicted by filled circles) were compared to a set of correlations obtained from 10,000 spin tests (depicted by box plots). Asterisks denote statistical significance (*p*_spin_ < 0.05, two-tailed). The association between node and neighbor atrophy was consistent across resolutions and significantly greater in empirical networks compared to null networks in both datasets. Right panel: Epicenter analysis was repeated using the binarized structural connectivity network. Epicenter likelihood rankings are depicted across the cortex. The most likely epicenters with high significant rankings are regions that are mainly located at the bilateral anterior insular cortex and temporal lobes (10,000 spin tests). The results are consistent with the original analysis where SC-defined neighbors were weighted by structural connectivity (Fig. 2).

**Figure S3.**
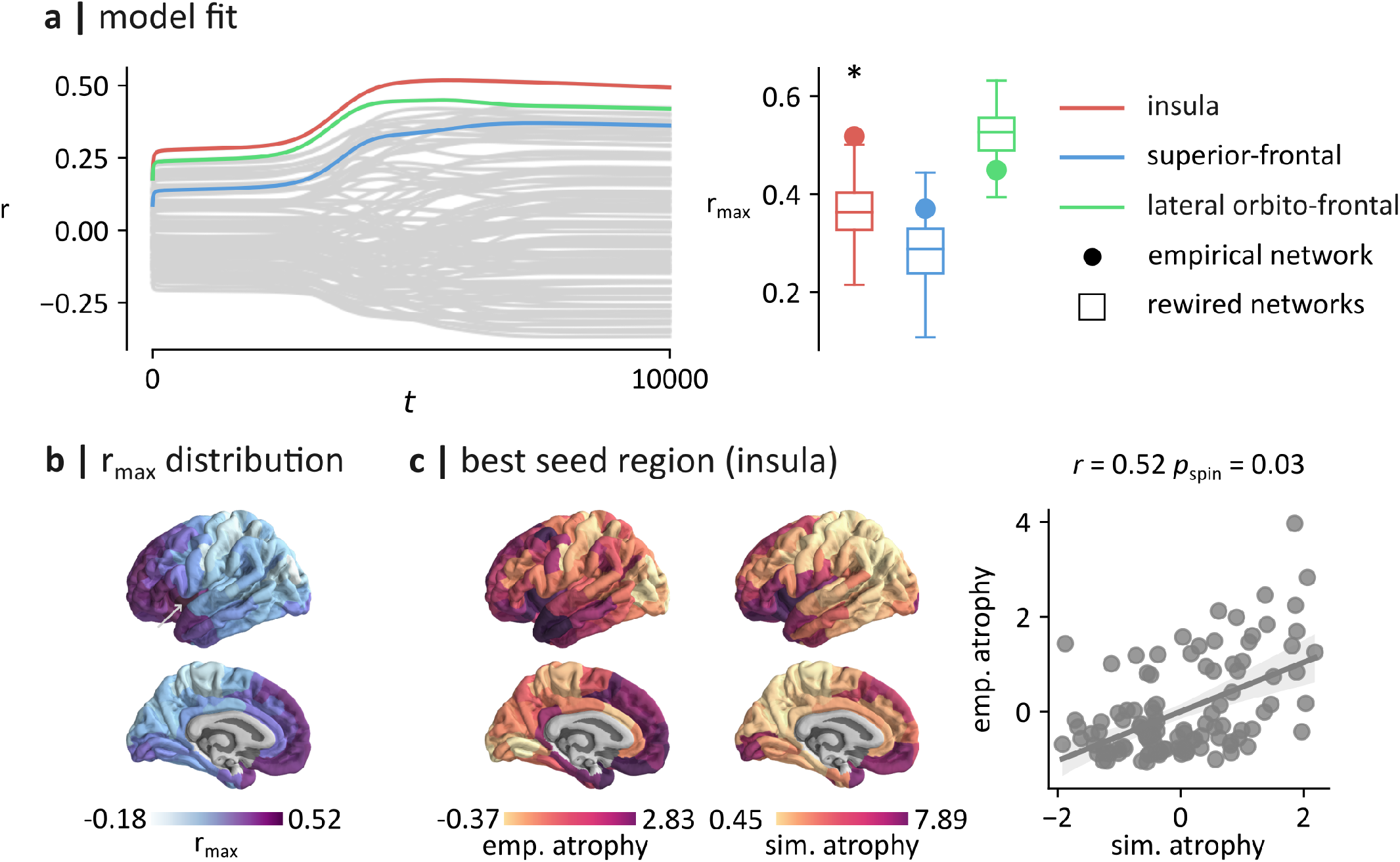
Simulation-based epicenter analysis using the GENFI dataset. (a) Left panel: the spreading process was initiated in every brain region and the correlation between the simulated and empirical patterns of atrophy was computed. The colored lines highlight the correlations obtained by seeding regions of the insula (*r*_max_ = 0.52; red), the superior-frontal cortex (*r*_max_ = 0.37; blue) and lateral orbito-frontal cortex (*r*_max_ = 0.45; green). The correlations for other brain regions are shown in gray. Right panel: to control for the potential effect of a brain region’s spatial embedding, *r*_max_ values were compared to *r*_max_ correlations obtained using rewired networks that preserve the wiring-cost of the empirical structural network. Asterisks denote statistical significance (*p* < 0.05, two-tailed). The *r*_max_ computed by seeding the insula of the empirical network (*r*_max_ = 0.52) was significantly larger than the *r*_max_ computed by seeding the insula of the rewired networks (*p* = 0.006). (c) The largest fit (*r*_max_) obtained by seeding each brain region is shown on the surface of the brain. Larger values of *r*_max_ were generally obtained by seeding insular and prefrontal regions. (d) Left panel: empirical pattern of atrophy (GENFI). Middle panel: simulated pattern of atrophy producing the maximal fit. This pattern of atrophy was obtained with the insula as the seed, and at *t* = 5730 (see the arrow in panel b). Right panel: scatter plot of the relationship between standardized empirical and simulated patterns of atrophy (*r* = 0.52, *p*_spin_ = 0.03).

## Appendix

### List of GENFI consortium members

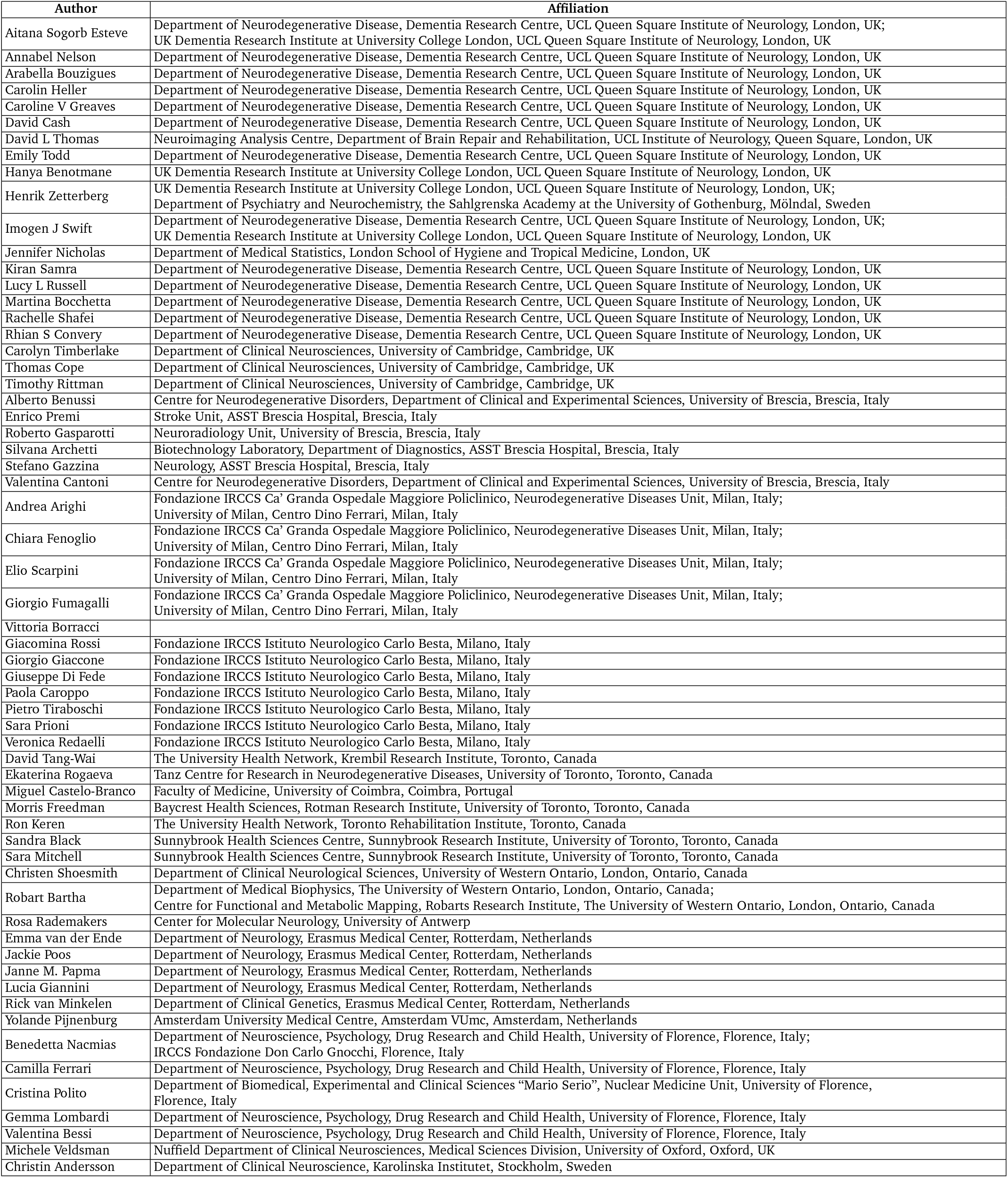
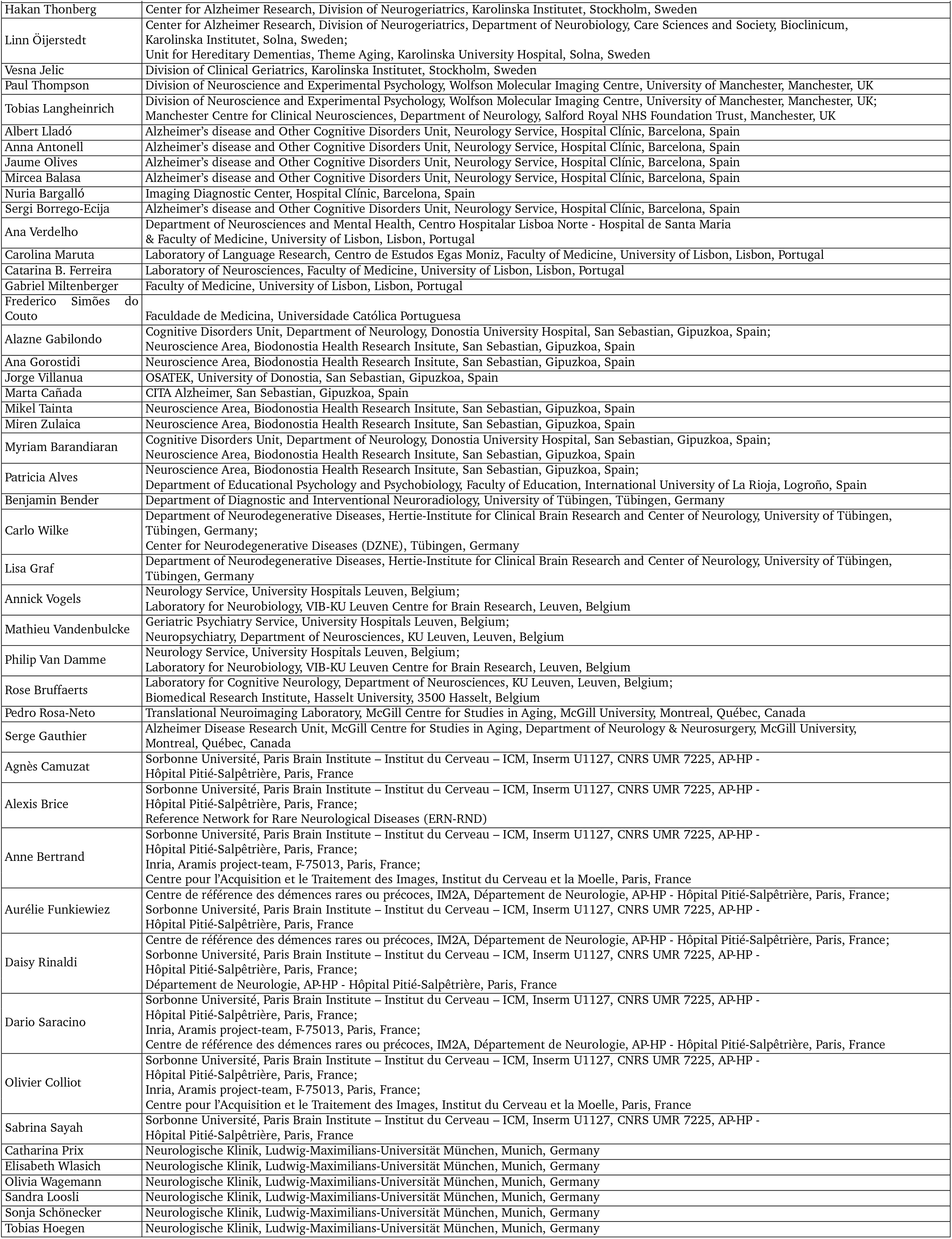
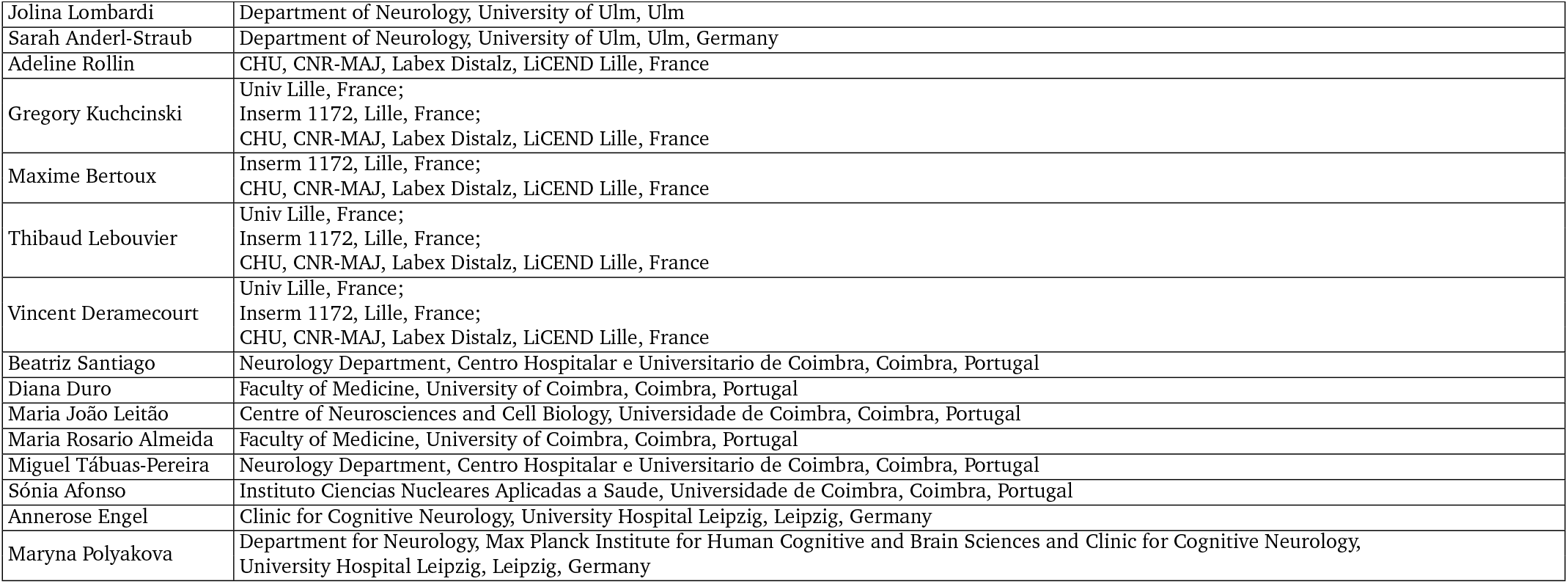

